# Free water in white matter differentiates MCI and AD from control subjects

**DOI:** 10.1101/537092

**Authors:** Matthieu Dumont, Maggie Roy, Pierre-Marc Jodoin, Felix C. Morency, Jean-Christophe Houde, Zhiyong Xie, Cici Bauer, Tarek A. Samad, Koene R. A. Van Dijk, James A. Goodman, Maxime Descoteaux, for the Alzheimers Disease Neuroimaging Initiative

## Abstract

Recent evidence show that neuroinflammation plays a role in many neurological diseases including mild cognitive impairment (MCI) and Alzheimer’s disease (AD), and that free water (FW) modeling from clinically acquired diffusion MRI (DTI-like acquisitions) can be sensitive to this phenomenon. This FW index measures the fraction of the diffusion signal explained by isotropically unconstrained water, as estimated from a bi-tensor model. In this study, we developed a simple FW processing pipeline that uses a safe white matter (WM) mask without gray matter (GM)/CSF partial volume contamination (*WM*_safe_) near ventricles and sulci. We investigated if FW inside the *WM*_safe_ mask, including and excluding areas of white matter damage such as white matter hyperintensities (WMHs) as shown on T2 FLAIR, computed across the whole white matter could be indicative of diagnostic grouping along the AD continuum.

After careful quality control, 81 cognitively normal controls (NC), 103 subjects with MCI and 42 with AD were selected from the ADNIGO and ADNI2 databases. We show that MCI and AD have significantly higher FW measures even after removing all partial volume contamination. We also show, for the first time, that when WMHs are removed from the masks, the significant results are maintained, which demonstrates that the FW measures are not just a byproduct of WMHs. Our new and simple FW measures can be used to increase our understanding of the role of inflammation-associated edema in AD and may aid in the differentiation of healthy subjects from MCI and AD patients.

## 1. Introduction

White matter (WM) atrophy in Alzheimer’s disease (AD) was observed more than three decades ago [1]. The microstrucutural changes observed in the WM of AD patients include axonal deterioration, Wallerian degeneration, loss of myelin density, loss of oligodendrocytes, microglia activation and vascualar degeneration [2, 3, 4, 5, 6, 7]. Numerous studies have shown that changes in the WM are an early event in the development of AD, happening in preclinical stages [3, 8, 9]. Changes in the microstructure of WM have even been reported before measurable hippocampal atrophy in mild cognitive impairment (MCI) [10] and preclinical AD [11]. More recent evidence shows that chronic neuroinflammation also contributes to the process of neurodegeneration in AD and was recently observed in the WM of AD patients [12].

Microglia-induced neuroinflamation in patients have been mostly studied using PET imaging ligands such as 11C-PK11195 [13]. However, to identify WM changes, diffusion MRI has been the modality of choice [14]. Studies, in the past decade, have identified various regions in the WM where the diffusion measures, mostly diffusion tensor imaging (DTI)-based measures such as fractional anisotropy and mean, axial, and radial diffusivities, correlate with symptoms of MCI and AD [15, 16, 17, 18, 19]. A more recent diffusion measure is the free water (FW) index, which measures the fraction of the diffusion signal explained by isotropically unconstrained water [20], as estimated from a regularized bi-tensor model. In white matter, this measurement represents either FW in extracellular space around axons or FW contamination from cerebrospinal fluid in adjacent voxels. An elevated FW index in white matter has been suggested to indicate neuroinflammation [21] and has been described in normal aging [22] and many neurological disorders such as schizophrenia [23, 24, 25], Parkinson’s disease [26], and AD [27, 28, 29].

In AD patients, an association between the widespread increased FW and poorer attention, executive functioning, visual construction, and motor performance was shown in [28, 29]. FW-corrected DTI measures are more sensitive to differentiate AD groups compared to standard DTI measures [29]. In a longitudinal study, FW-corrected radial diffusivity, but not un-corrected radial diffusivity, was higher in the WM of MCI patients who converted to AD compared to MCI patients who did not convert [27]. FW-corrected DTI measures also demonstrate greater sensitivity to associations between AD pathology and white matter microstructure compared to standard DTI measures [11].

In this study, we developed a FW processing pipeline that uses a WM “*safe*” mask (*WM*_safe_) without GM/CSF partial volume contamination and computed new and simple whole-brain FW measures based on this *WM*_safe_ mask (including and excluding WMHs within the mask) for three different groups (cognitively normal, MCI and AD subjects), selected from the ADNIGO and ADNI2 databases. The prevalence and degree of WMHs is known to increase with age [30] and therefore WMHs cannot be ignored in WM processing of aged groups. We show that our FW measures were significantly higher in MCI and AD groups compared to NC, but only when using a WM safe mask. We also show, for the first time, that when WMHs are removed from the mask, the significant results remained, which demonstrates that FW measures are not just a byproduct of WMHs.

## 2. Methods

### 2.1. Study participants

226 subjects from the ADNIGO and ADNI2 databases passed the necessary quality assurance (QA) phases of the diffusion MRI analysis pipeline (described below). Of those participants, 81 (38 males, 43 females) were cognitively normal (normal control, NC), 103 (69 males, 34 females) had a diagnosis of mild cognitive impairment (MCI) and 42 (25 males, 17 females) had a diagnosis of AD. Age range per group was between 67 and 95 years for NCs, between 60 and 95 years for MCIs and bewteen 61 and 97 for ADs. Mean age was 78.46 for NCs, 79.0 for MCIs and 79.38 for ADs. All participants had good general health, good hearing and seeing abilities, no depression or bipolar condition, no history of alcohol or drug abuse and completed at least six grades of education. Also, NCs had no memory impairment and their clinical dementia rating (CDR) was 0. MCI subjects included early and late MCI with impaired memory and a CDR of 0.5, while AD subjects met criteria for dementia and had a CDR between 0.5 and 1 [31]. Participants did not suffer from any neurological disorders other than MCI and AD such as brain tumor, multiple sclerosis, Parkinson’s disease, or traumatic brain injury.

### 2.2. MRI data acquisition

Data used in the preparation of this article were obtained from the Alzheimers Disease Neuroimaging Initiative (ADNI) database (adni.loni.usc.edu). The ADNI was launched in 2003 as a public-private partnership, led by Principal Investigator Michael W. Weiner, MD. The primary goal of ADNI has been to test whether serial magnetic resonance imaging (MRI), positron emission tomography (PET), other biological markers, and clinical and neuropsychological assessment can be combined to measure the progression of mild cognitive impairment (MCI) and early Alzheimers disease (AD). For up-to-date information, see www.adni-info.org.

Of the available modalities, we used the T1w, diffusion weighted imaging (DWI) and fluid attenuation inversion recovery (FLAIR) scans. The DWI scans were acquired along 41 evenly distributed directions using a b-value of 1000 s/mm^2^ with a 1.3 × 1.3 × 2.7 mm^3^ spatial resolution. The T1w and FLAIR scans were acquired at 1.2 × 1.05 × 1.05 and 0.85 × 0.85 × 5 mm^3^ spatial resolution, respectively. Data was acquired at 58 different North-American locations.

### 2.3. MRI processing pipeline

The processing pipeline is illustrated in Figure 1. At first, the T1w and DW images were denoised with a non-local means method robust to Rician noise [32], followed by an MRI bias field correction performed with ANTs N4 correction tool [33]. The brain mask (BM) was then processed and the skull was removed using the BEaST brain extraction software [34]. We referred to these methods as the *preprocessing* step in Figure 1. Then, the T1w and FLAIR images were nonlinearly registered to the 1×1×1 up-sampled diffusion space with ANTs registration [33]. Tissue segmentation was then performed on the transformed T1w to obtain a binary map of the CSF, GM, and WM. This was done using ANTs Atropos [33]. In order to prevent any CSF contamination in regions susceptible to partial volume effect, a “safe WM mask” (*WM*_safe_) was built by combining the following morphological operations on the CSF, WM, GM and brain binary masks:

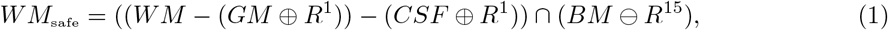

where *R^n^* is a 3D structuring element of radius *n*, ⊕ is the dilatation operator, ⊖ is the erosion operator and ∩ the intersection operator as illustrated in Figure 1. Using the FLAIR and T1w images, a binary map of WMHs was also computed using *volBrain* [35]. The bi-tensor model proposed by Pasternak et al. [20] was fit onto the DW signal. The result of this fit is a fraction representing the contribution of unconstrained water to the original signal and a new signal representing the tissue contribution. The fraction of unconstrained water contribution in a voxel is what we commonly call FW volume and the 3D image of this FW volume is called the FW map. The tissue signal is the FW-corrected DWI signal, as it represents the signal without its unconstrained component. The safe white matter mask, the WMH mask, and the FW map were then used to extract the mean FW value (*μFW*) and the relative FW volume (*rFW*). The *rFW* is the total volume of FW voxels within the safe white matter mask with FW values greater than 0.1, divided by the total volume of the safe white matter mask. *rFW_m_* and *μFW_m_* are defined as such:

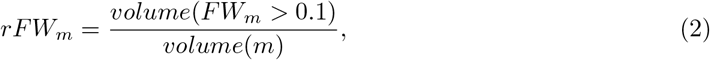

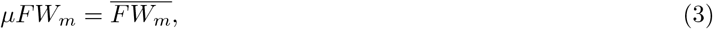

where *m* ∈ {*WM*_safe_, *WMHs*, *WM*_safe_ – *WMHs*}. The 0.1 threshold was chosen to minimize the impact of noise on the rFW metric. All processing was done using a Nextflow [36] pipeline with all software dependencies bundled in a Singularity container [37] ensuring quick and easy reproducibility of the results.

**Figure 1:**
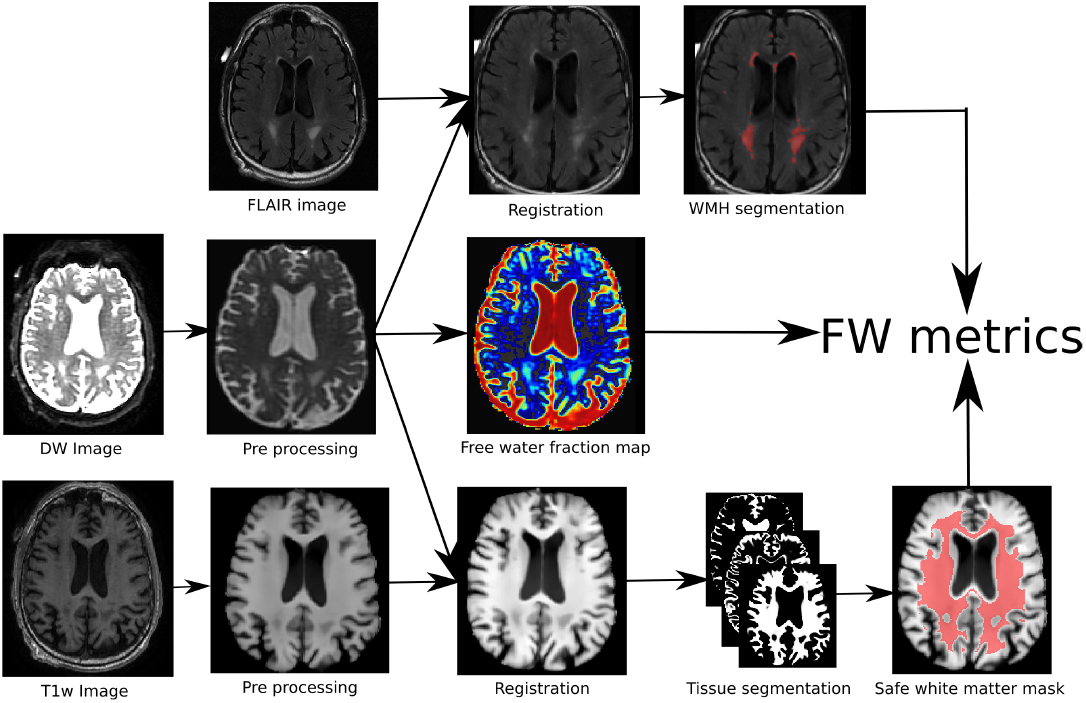
Pipeline of the proposed method: (1) the DWI and T1w images are first preprocessed, (2) the three modalities are co-registered of which (3) are extracted the FW map, the tissue map and the WMHs areas. (4) the combination of the three maps leads to the proposed FW metrics.

### 2.4. Statistics

A cross-sectional analysis was performed at the first available time point comparing rFW and *μ*FW in NC (n=81), MCI (n=103), and AD (n=42). An analysis of variance (ANOVA) was performed to test for a main effect of diagnostic group followed by a post-hoc pairwise Tukey test to assess differences between sub groups [38]. A log transformation was applied to the rFW and *μ*FW metrics to improve normality of the distribution before analyses [39].

### 2.5. Quality assurance

Out of all available subjects in ADNI2 and ADNIGO, 239 had at least one time point with all the images required (T1w, DWI and FLAIR) to go through the processing pipeline. Visual QA was performed on all images of all time points and those with problems impossible to correct (missing brain parts, acquisition artifacts) were rejected. Gradient information was also QA-ed to make sure every DWI image had 41 evenly distributed direction on one single acquisition shell. This first QA pass eliminated 9 subjects bringing the count of subjects with usable data to 230. Visual inspection was performed on brain extraction of T1w and DWI as well as on the non-linear registration of the FLAIR on the T1w and of the T1w on DWI. Every tissue segmentation mask (WM, GM, CSF) as well as the WMH mask was inspected. This second QA pass eliminated 4 subjects, 3 with artefacts in the DWI images causing improbable values in metrics and one with an obviously incorrect T1 brain mask, leaving 226 subjects with usable data for the group analysis.

## 3. Results

As shown in Table 1 results of the initial ANOVA tests show a significant main effect of group membership across all regions of interests. Post-hoc Tukey tests show that both *rFW* and *μFW* are significantly higher in the *WM*_safe_ for MCI and AD subjects than for NC subjects whether or not *WMHs* were included. Interestingly, when looking at *rFW* and *μFW* specifically within the WMH mask we do see significant between-group differences but with lesser effect and neither of them being able to separate both NC-MCI and NC-AD.

**Table 1:** The F-statistic obtained from the ANOVA test is displayed in the first column and the rest of the table shows the Tukey post-hoc pairwise group differences(on log-scale) with the standard error in parentheses. The statistical significance(in bold) is shown as: * p<0.05, ** p<0.01, *** p<0.001.

|  | F(2, 223) | NC-MCI(SE) | NC-AD(SE) | MCI-AD(SE) |
| --- | --- | --- | --- | --- |
| $rFW_{WM_{safe}-WMHs}$ | **16.02\*\*\*** | **-0.52(0.18)\*\*\*** | **-0.86(0.23)\*\*\*** | -0.34(0.20) |
| $\mu FW_{WM_{safe}-WMHs}$ | **13.58\*\*\*** | **-0.49(0.17)\*\*\*** | **-0.70(0.19)\*\*\*** | -0.21(0.20) |
| $rFW_{WM_{safe}}$ | **12.79\*\*\*** | **-0.53(0.19)\*\*\*** | **-0.90(0.22)\*\*\*** | -0.36(0.21) |
| $\mu FW_{WM_{safe}}$ | **12.66\*\*\*** | **-0.50(0.17)\*\*\*** | **-0.74(0.20)\*\*\*** | -0.23(0.20) |
| $rFW_{WMHs}$ | **4.14\*** | **-0.13(0.07)\*** | -0.14(0.10) | -0.01(0.09) |
| $\mu FW_{WMHs}$ | **3.80\*** | -0.13(0.08) | **-0.18(0.10)\*** | -0.05(0.10) |

To represent the spatial differences in free water repartition between groups, every T1W image already registered in diffusion space was non-linearly registered to the MNI152 space with the ANTs registration tool [33]. The resulting transformations were applied to the free water volumes. Mean and standard deviation free water volume for each group was computed and used to obtain a z-score volume of each subject compared to each group. These z-score volumes were averaged and thresholded at z ≥ 2 standard deviations to obtain binary group comparison volumes.

In both the NC vs AD and NC vs MCI comparisons, the voxels showing differences are mostly located in the corticospinal tract (CST) and bundles of the limbic system such as the cingulum and the fornix. Figure 2 shows that intensity and location of significant z-score clusters is different when comparing AD or MCI to NC.

**Figure 2:**
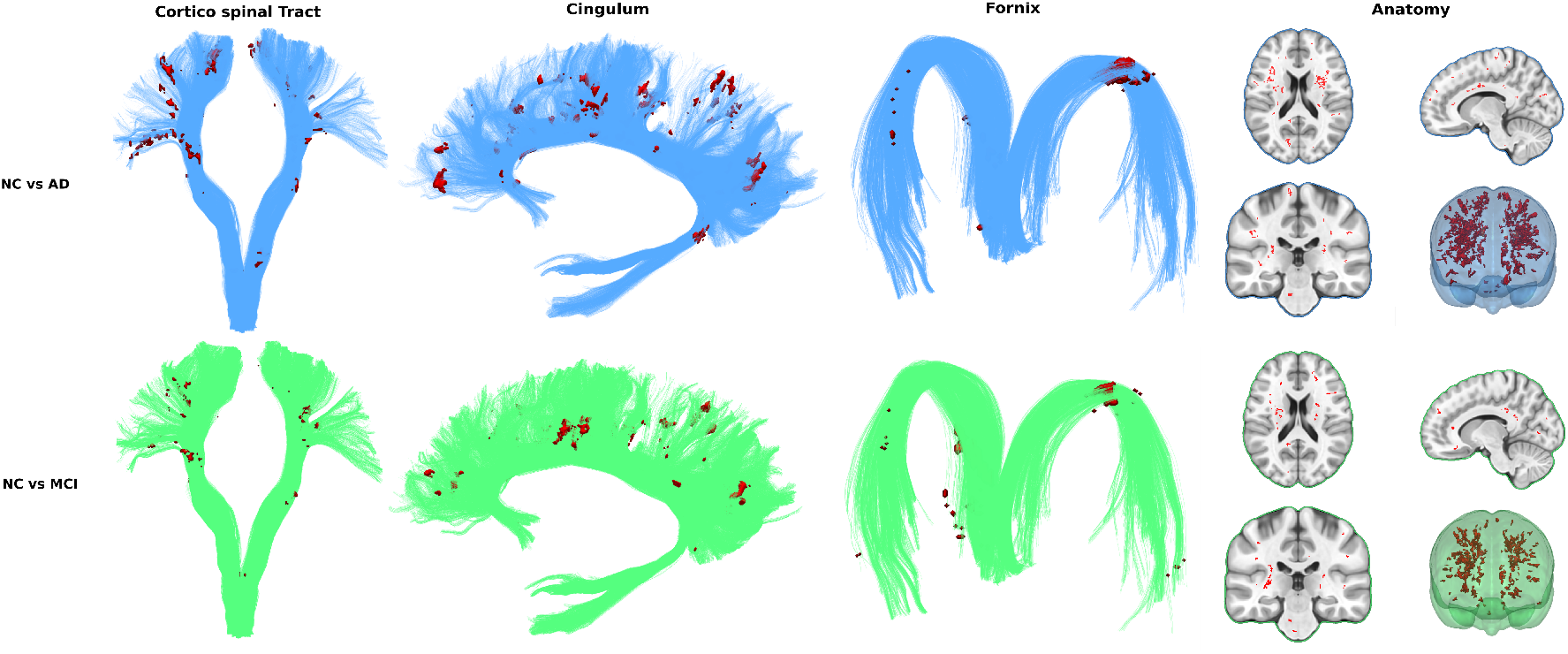
Spatial repartition in free water differences across groups

## 4. Discussion

We have shown that MCIs and ADs have significantly higher FW metrics compared to NCs. There was a correlation between FW and age but it was found to be weak and consistent across all groups. We also demonstrated that the use of a safe WM mask is critically important to prevent partial volume contamination that can perturb the overall results. Our new and simple FW measures can be used to increase our understanding of the role of inflammation-associated edema in AD and may aid in the differentiation of healthy subjects from MCI and AD patients. Significant results were also obtained when WMHs were removed from the WM mask, which demonstrate that FW metrics differences between group were not due to white matter lesions that can be seen on T2 FLAIR images as WMHs. In fact, a key observation of this study is that FW in MCI and AD subjects is globally distributed through the WM and is not specifically associated to WMHs or other regions typically associated to cognition like the limbic system.

Neither FW metrics could differentiate between MCI and AD subjects. This result could be attributed to the study design since measuring FW metrics in non-specific anatomical brain region such as the whole WM gives a very broad picture of the FW content in the brain, which may hide localized differences. Analyzing FW content along specific WM bundles would be expected to yield more specific results. To do so, tractography would then be used to reconstruct the global WM architecture followed by an automated segmentation of several key WM bundles such as the fornix, cingulum, corpus callosum, and association tracts (arcuate fasciculus, uncinate, inferior longitudinal and inferior fronto-occipital fasciculus). FW metrics would be analyzed along those bundles, as done in apparent fiber quantification (AFQ) [40] and tract-profiling [41]. Future work would also include looking at how FW correlates with abeta and tau data available in ADNI to further support the hypothesis that FW is a good correlate measurement of neuroinflammation.

However, FW metrics also have limitations, i.e. they are derived from a bi-tensor model, which is limited to representing a FW compartment and a single fiber population. However, it is estimated that 66 to 90 percent of brain WM voxels contain at least two fiber populations [27, 28]. In those voxels, the estimated contribution of the FW compartment is incorrectly estimated, since some signal arising from the fiber populations not fitted to the single fiber tensor will be assigned to the FW compartment. To correct this bias, a FW model accounting for more than one fiber population would need to be used to better fit the signal. While a more sophisticated model would certainly better characterize the information contained in the non-free-water portion of the signal and give more accurate free water indices, these models require multi-shell DWI acquisitions which are unavailable in ADN2 and ADNIGO.

## 5. Conclusion

This study demonstrates that after removing partial volume contamination as well as WMHs lesions, the free water content of healthy looking white matter differentiates MCI and AD groups from healthy subjects. Our method is based on existing DTI-like diffusion data, is atlas free, requires no registration with a reference brain, no PET scan, no tractography, has few tunable parameters, and takes a few minutes only of computation. The method is a simple but powerful approach that may be used in the context of patient selection and stratification for novel treatments that are aimed at treating or preventing inflammation components of AD using legacy or standard diffusion MRI data. The significant differences of our FW metrics between NC and MCI as well as NC and AD may demonstrate the potential of FW as a tool to study neuroinflammation. We intend to extend this work with analyses of FW metrics in specific white matter bundles and sections of bundles. Also, characterization over time of our new FW metrics in an MCI population could help differentiate those older adults who will remain relatively stable and those who will progress to AD, which has utility for patient selection and stratification of subjects in preclinical stages of AD.

## 6. Acknowledgements

This research has been evenly funded by Imeka Solution inc. and Pfizer inc.

Data collection and sharing for this project was funded by the Alzheimer’s Disease Neuroimaging Initiative (ADNI) (National Institutes of Health Grant U01 AG024904) and DOD ADNI (Department of Defense award number W81XWH-12-2-0012). ADNI is funded by the National Institute on Aging, the National Institute of Biomedical Imaging and Bioengineering, and through generous contributions from the following: AbbVie, Alzheimers Association; Alzheimers Drug Discovery Foundation; Araclon Biotech; BioClinica, Inc.; Biogen; Bristol-Myers Squibb Company; CereSpir, Inc.; Cogstate; Eisai Inc.; Elan Pharmaceuticals, Inc.; Eli Lilly and Company; EuroImmun; F. Hoffmann-La Roche Ltd and its affiliated company Genentech, Inc.; Fujirebio; GE Healthcare; IXICO Ltd.; Janssen Alzheimer Immunotherapy Research & Development, LLC.; Johnson & Johnson Pharmaceutical Research & Development LLC.; Lumosity; Lundbeck; Merck & Co., Inc.; Meso Scale Diagnostics, LLC.; NeuroRx Research; Neurotrack Technologies; Novartis Pharmaceuticals Corporation; Pfizer Inc.; Piramal Imaging; Servier; Takeda Pharmaceutical Company; and Transition Therapeutics. The Canadian Institutes of Health Research is providing funds to support ADNI clinical sites in Canada. Private sector contributions are facilitated by the Foundation for the National Institutes of Health (www.fnih.org). The grantee organization is the Northern California Institute for Research and Education, and the study is coordinated by the Alzheimers Therapeutic Research Institute at the University of Southern California. ADNI data are disseminated by the Laboratory for Neuro Imaging at the University of Southern California.

